# Heparanase 2 regulates endothelial permeability and prevents from proteinuria via VEGFA and FGF signaling

**DOI:** 10.1101/2024.08.14.608012

**Authors:** Yannic Becker, Sergey Tkachuk, Anne Jörns, Ahmad Alwakaa, Jan Hegermann, Heiko Schenk, Yulia Kiyan, Hermann Haller

## Abstract

**Background:** Heparan sulfates (HS) attached to the apical surface of vascular endothelial cells (ECs) play an important role in regulating endothelial permeability and ligand recognition by cell-surface receptors. Shedding of heparan sulfate (HS) from the EC surface increases vascular leakage and is associated with vascular diseases. Recently, heparanase 2 (Hpa2) was described as a novel regulatory molecule that controls HS shedding. However, its role in regulating HS physiology in the vascular endothelium is largely unknown. Here, we characterize the role of endogenous Hpa2 in the vertebrate vascular system.

**Methods:** We use zebrafish larvae as our primary animal model. Hpa2 expression and localization was examined by *in situ* hybridization and immunofluorescence. Hpa2 loss-of-function (LOF) was induced by CRISPR-Cas9 or morpholino antisense strategies. We assessed vascular permeability, blood vessel architecture, and EC morphology using transgenic zebrafish and transmission electron microscopy. EC expression profiles and HS quantity were analyzed in Hpa2-LOF larvae. The capacity of recombinant Hpa2 to modulate signaling in ECs by the HS-binding growth factors fibroblast growth factor 2 (FGF2) and vascular endothelial growth factor A_165_ (VEGFA_165_) was tested by western blotting and immunofluorescence. Attenuation of the Hpa2-LOF phenotype was tested *in vivo* in zebrafish larvae via use of recombinant Hpa2 and pharmacological inhibition of FGF and VEGFA signaling.

**Results:** We detected *hpse2* expression in hepatic tissue and localized the protein in blood vessels. Hpa2-LOF larvae exhibited increased vascular permeability, occasional hypersprouting, and altered EC and extracellular matrix (ECM) morphology. Hpa2-LOF also reduced HS levels and caused changes in the endothelial transcriptome characterized by dysregulated genes involved in ECM-receptor interaction and signal transduction regulation. Recombinant hHpa2 rescued the Hpa2-LOF phenotype in zebrafish. We showed *in vitro* that Hpa2 competes with FGF2 and VEGFA_165_ for binding on the EC surface and consequently reduces the cellular response these factors elicit. Pharmacological inhibition of these pathways alleviated the Hpa2*-*LOF phenotype in zebrafish.

**Conclusion:** We conclude that Hpa2 is a circulating molecule that maintains vascular integrity by regulating HS-dependent processes on the EC surface. These results may translate into novel strategies applying recombinant Hpa2 to treat microvascular diseases.

## Introduction

Endothelial cells (ECs) form the inner layer of blood vessels and thus are critically important for systemic function of multi-organ species. To adapt to different organ-specific conditions and changes in the environment, ECs receive a broad range of signals including mechanical, cell-cell contact, and paracrine and endocrine inputs^1^. Many of these signals are sent via the bloodstream, thus stimulating ECs from the luminal side, where ECs are covered by a pericellular matrix termed the endothelial glycocalyx (eGCX). The eGCX is a carbohydrate-rich, highly dynamic layer that is closely linked to vascular physiology and disease^2,3^. It participates in diverse functions including sensing of shear stress^4^, and regulation of vascular permeability^5^, platelet adhesion and coagulation^6^, leukocyte adhesion and inflammation^7^, and overall vascular homeostasis^8^.

The eGCX consists primarily of membrane-bound proteoglycans, which are core proteins attached to one or more long linear polysaccharides termed glycosaminoglycans (GAGs); and plasma proteins, including growth factors, complement components, and coagulation proteins, adhering to GAG chains. The most abundant GAG is heparan sulfate (HS), and HS proteoglycans are enriched in the eGCX, playing major roles in both vascular function and disease. HS, which comprises repeating disaccharide units of N-acetylglucosamine and hexuronic acid, undergoes extensive chain modifications during synthesis in the Golgi compartment including deacetylation, sulfation, and epimerization, yielding diverse structural motifs within HS chains^9^. This structural diversity provides specific binding sites for interaction partners, which has led to the discovery of more than 300 proteins that interact with HS^10^. Among them are growth factors essential for endothelial physiology, including members of the fibroblast growth factor (FGF) and vascular endothelial growth factor (VEGFA) family^1^.

Degradation of the eGCX, including HS, is closely associated with microvascular diseases including sepsis^11^, ischemia-reperfusion injury^12^, and diabetic microangiopathy^13^ all of which leave the endothelium in an inflammatory, thrombotic, and eventually dysfunctional state^14^. Major contributors to eGCX degradation include glycosidases, which cleave and release GAG chains from core proteins. The endo-β-D-glucuronidase heparanase 1 (Hpa1) is the only enzyme in mammals known to be capable of degrading HS from the EC surface^15^. Hpa1 is upregulated in cardiovascular diseases including diabetes^16^, sepsis^11^, and atherosclerosis^17^.

Shortly after the discovery of Hpa1 in 1999, McKenzie and colleagues cloned and characterized a homolog of Hpa1, which they named heparanase 2 (Hpa2), that shares up to 40% sequence homology^18^. Importantly, Hpa2 possesses no catalytic activity towards HS. Instead, Hpa2 binds HS with strong affinity, which led to the model that Hpa2 competes with Hpa1 for HS substrate binding, thereby blocking Hpa1-mediated HS degradation^19^. To date, the physiological function of Hpa2 has been investigated primarily in the context of cancer biology, where it has been shown to be a counterpart of the tumor-promoting, pro-metastatic Hpa1, although this role depends on the tumor subtype ^20,21^. In contrast, little is known about the role of Hpa2 in eGCX and EC physiology. Recently it was shown in sepsis models that Hpa2 overexpression in ECs protects from Hpa1-mediated eGCX degradation and alleviates pro-inflammatory signaling^22^. In addition, intravenous application of Hpa2 and Hpa2-derived peptides alleviated kidney dysfunction in a lipopolysaccharide-induced glomerulonephritis model^23^. However, the endogenous function of Hpa2 in the vascular system *in vivo* has not yet been characterized, and the mechanism through which it may protect the endothelium is largely unknown.

The zebrafish *Danio rerio* is an excellent model organism for addressing questions about Hpa2 and the eGCX. Zebrafish develop rapidly, providing a functional circulatory system within 24 hours post fertilization (hpf) that can be monitored in detail due to larvae transparency^24^. Using zebrafish larvae, we provide data that Hpa2 circulates in the vascular system of vertebrates, where it is necessary to maintain EC integrity and vascular homeostasis. In addition, we demonstrate structural and functional conservation of Hpa2 among vertebrates, thereby highlighting the translational potential of our findings. Finally, we show that Hpa2 regulates HS-dependent growth-factor signaling of fibroblast growth factor 2 (FGF2) and vascular endothelial growth factor A_165_ (VEGFA_165_) through its strong binding affinity towards HS. We postulate that Hpa2 has a systemic regulatory function in the endothelium by balancing HS-dependent signal input for ECs.

## Results

### Hpa2 is a conserved protein that circulates in the vasculature of vertebrates

Hpa2 has been characterized primarily in view of its roles in normal development of the peripheral nervous system and in neoplasias, deploying human- and murine-derived samples as primary research material^25,26^. Prompted by recent findings that exogenous Hpa2 may have protective effects for the endothelium^22,23^, here we aimed to elucidate the role of endogenous Hpa2 in the vertebrate vascular system. First, we compared the sequence of Hpa2 from higher vertebrates with that of Hpa2 from *Danio rerio* and found high sequence conservation at the protein level, with 72% identity and 83% similarity (Figure 1A, B). The C-terminal region of the protein, which includes the putative HS-binding domains, is particularly highly conserved (Figure 1B). Concordantly, modeling and alignment of human and zebrafish Hpa2 revealed structural conservation (Figure 1C).

**Figure 1:**
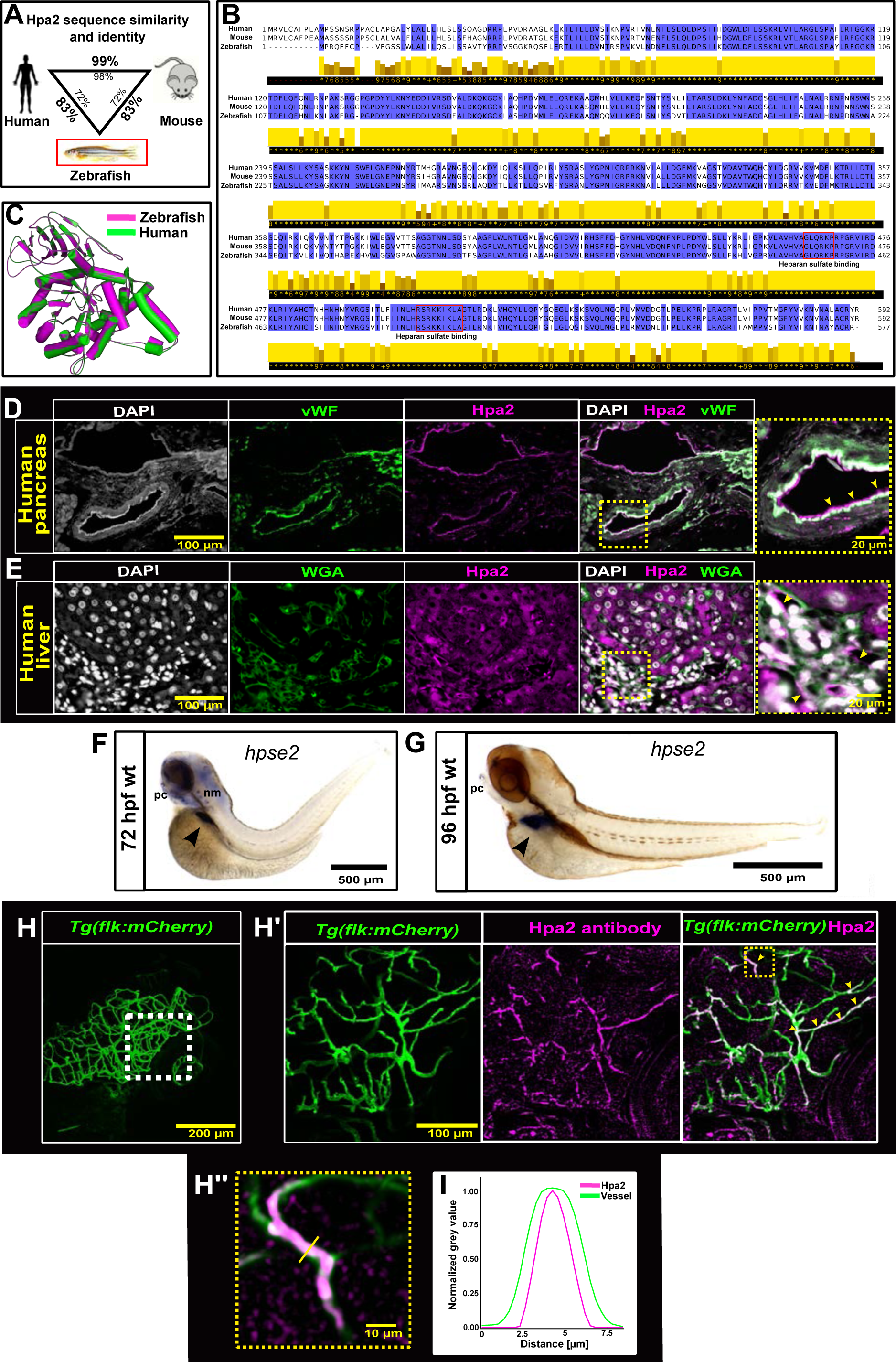
Heparanase 2 (Hpa2) is structurally conserved in vertebrates, localizes on the luminal side of the vasculature, and is expressed by hepatic tissue in zebrafish. **(A)** Hpa2 protein sequence similarity and identity between humans, mice, and zebrafish. Percentages outside the triangle refer to percentage identity, percentages inside to similarity. **(B)** Multiple sequence alignment of Hpa2. Conserved residues marked in blue; amino-acid similarity is scored 1 (low) to 9 (high) below the sequence. Putative heparin-/heparan-sulfate-binding sequences underlined in red. **(C)** Predicted protein structure and spatial alignment of human (green) and zebrafish (magenta) Hpa2. **(D,E)** Immunofluorescence staining of human pancreatic **(D)** and liver parenchyma (E). **(D)** Pancreatic tissue stained for Von Willebrand Factor (vWF; an EC marker) and Hpa2. Region of interest (ROI) magnifies vWF-rendered endothelium with luminal signal for Hpa2 (yellow arrowheads). **(E)** Hepatic tissue stained for wheat germ agglutinin (WGA) and Hpa2. ROI indicates Hpa2 signal in hepatic tissue and capillaries (yellow arrowheads). **(F,G)** Whole-mount *in situ* hybridization for *hpse2* in wildtype larvae at 72 hours post-fertilization (hpf) (**F**) and 96 hpf (**G**). Arrowhead points towards hepatic tissue. pc = pharyngeal cartilage, nm = neuromasts. **(H)** Whole-mount immunofluorescence for Hpa2 in vasculature-labeling *Tg(flk:mCherry)* zebrafish at 96 hpf. Image shows z-projection of larval cranial vasculature. (**H’,H’’**) ROI of *Tg(flk:mCherry)* fish stained for Hpa2. Arrowheads indicate colocalization of Hpa2 antibody signal and mCherry signal. (**I**) Signal intensity plots of mCherry and Hpa2 antibody across blood vessel normalized to the maximum grey value in each channel.

We showed previously that Hpa2 circulates in human plasma^27,28^. To further investigate the localization and origin of circulating Hpa2, we stained hepatic and pancreatic tissue samples from humans and mice for Hpa2 (Figure 1D, E, Figure S1). We detected Hpa2 on the luminal side of both small and large vessels in pancreatic tissue (Figure 1D, Figure S1B) and on the luminal side of hepatic sinusoids (Figure 1E, Figure S1C). Interestingly, Von Willebrand factor-positive ECs did not show a strong signal for Hpa2 (Figure 1D, Figure S1B). In contrast, the organ parenchyma of liver and pancreas expressed Hpa2 (Figure 1E, Figure S1C). Pancreatic beta cells were negative for Hpa2 (Figure S1A).

Using zebrafish as our primary research model, we first asked whether Hpa2 is expressed in the same organs in zebrafish as those where we observed expression in mouse and human. Whole-mount *in situ* hybridization detected Hpa2 expression in hepatic tissue of zebrafish larvae by purple color reaction starting at 72 hpf (Figure 1F, G). Specific expression was also detected in neuromast populations at 72 hpf (Figure 1F) and pharyngeal cartilage at 96 hpf (Figure 1G). After identifying the liver as the primary organ of *hpse2* expression in the larval stage, we applied the same antibody that we used to localize Hpa2 protein in murine and human tissue. In the EC-labeling *Tg(kdrl:mCherry)* zebrafish line, we detected co-localization of the Hpa2 antibody with zebrafish vasculature (Pearson’s coefficient = 0.56) (Figure 1H). Intensity profiles for the Hpa2 antibody were narrower than the mCherry signal derived from the EC (Figure 1I), suggesting that Hpa2 localizes within blood vessels. Because *hpse2* is expressed in neuronal tissue in other species^26,29^, we performed *in situ* hybridizations for *hpse2* in earlier developmental stages and detected expression at 24 hpf in the lateral line primordium, dorsal root ganglia, and pectoral fin buds (Figure S2A, B). At 48 hpf, expression was no longer detected along the spinal cord, but specific staining was observed in the pectoral fin tip as well as in neuromasts in the inner ear region (Figure S2C). Together, these data emphasize the conservation of Hpa2 protein structure and expression in vertebrates and provide a basis for further investigation of the vascular role of Hpa2 using zebrafish as a model system.

### Hpa2 loss-of-function (LOF) increases vascular permeability in zebrafish

To investigate the role of Hpa2 in the zebrafish vasculature, we used CRISPR-Cas9 technology to generate F0 Hpa2-LOF larvae. We targeted exon 1 and exon 5 of the *hpse2* gene using two different guide RNAs (one for each exon), and validated successful mutagenesis by fragment analysis of the *hpse2* locus from individual larvae (Figure S3A). Compared to control-injected embryos, the survival rate of mutated embryos decreased by 25% within the first two days of development (Figure S3B). To date, we have been unable to breed homozygous *hpse2* mutants, which implies embryonic lethality in *hpse2^-/-^* larvae. CRISPR-Cas9-induced Hpa2 LOF caused a pericardial edema phenotype in 30% (guide1) and 16% (guide2) of the surviving larvae (Figure 2A, B, C), implying a cardiovascular role for Hpa2. To further investigate the phenotype, we used *Tg(l-fabp:DBP-EGFP)* larvae, which secrete a 78-kDa fluorescently labeled vitamin-D-binding protein (VDB) into the vascular system^30^. Loss of fluorescence can be quantified from the retinal vessel plexus and serves as a surrogate marker for loss of plasma protein, indicating increased vascular permeability (Figure 2D). CRISPR-Cas9-induced Hpa2 LOF caused a 30% decrease in maximum eye fluorescence (MEF) for guide 1 and 32% for guide 2 (Figure 2E, F). To validate the effects of the CRISPR system for *hpse2,* we designed a splice-interfering morpholino (MO) for the intron-exon junction 10 of the *hpse2* pre-mRNA. Splice interference of the MO was confirmed by reverse transcriptase polymerase chain reaction (Figure 2H). The MO caused no specific edema phenotype and no decrease in survival relative to control embryos (Figure 2G, Figure S3B). MO-induced Hpa2 LOF induced a 12% loss in MEF using 0.58 pmol MO per embryo and 30% loss in MEF using 1.15 pmol per embryo (Figure 2J). Lower concentrations of MO failed to induce a consistent vascular-leakage phenotype (data not shown). Co-injection of human *hpse2* mRNA together with the *hpse2* MO rescued the Hpa2-LOF phenotype (Figure 2K). As an additional method to validate the vascular-leakage phenotype in Hpa2-LOF larvae, we injected 70 kDa fluorescently labeled dextran into the vasculature and observed a 55% increase in dextran leakage from Hpa2-LOF compared to control larvae, confirming the increased vascular permeability induced by Hpa2 LOF (Figure S3C).

**Figure 2:**
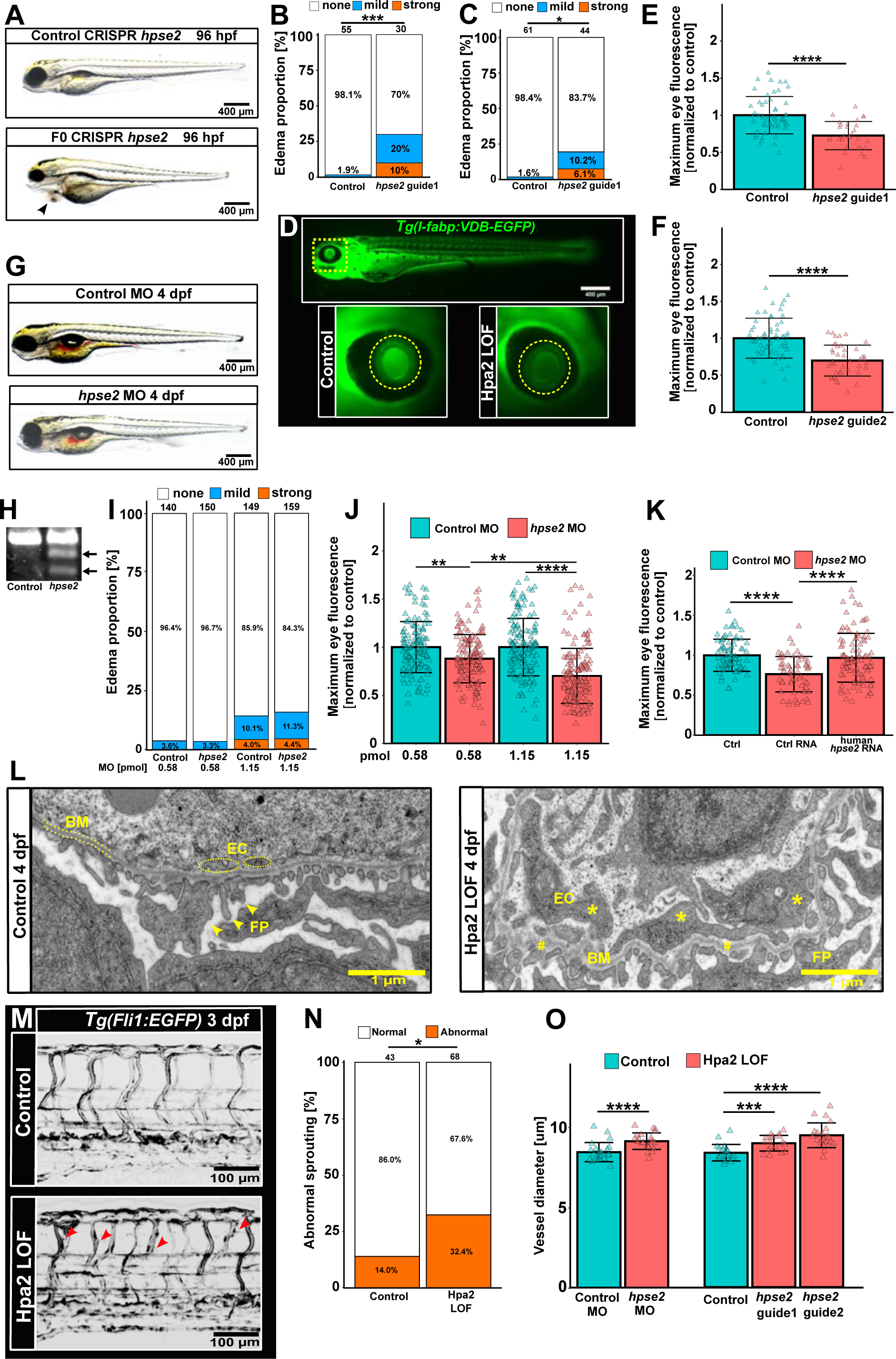
Hpa2 loss-of-function (LOF) induces vascular hyperpermeability and morphological abnormalities of endothelial cells (ECs) in zebrafish. (**A**) Lateral views of control and CRISPR-Cas9-induced F0 Hpa2-LOF larvae at 96 hpf. Arrowhead indicates pericardial edema formation. (**B,C**) Edema scoring of Hpa2-LOF larvae induced by two different guide RNAs targeting exon 1 (B) and exon 5 (C) of *hpse2*. The numbers above the columns indicate the number of examined larvae. **(D)** Use of *Tg(l-fabp:VDB-eGFP)* larvae to study vascular integrity in Hpa2-LOF larvae. Fish express liver-derived, EGFP-labeled vitamin-D-binding protein (VDB) as a fluorescent tracer that circulates in the vascular system. Fluorescence loss is quantified from the retinal vessel plexus (yellow circle). (**E,F**) Quantification of vascular permeability in CRISPR-induced Hpa2-LOF larvae at 96 hpf. Data points refer to individual larvae. **(G)** Lateral views of control and morpholino (MO)-induced Hpa2-LOF larvae at 96 hpf. (H) *hpse2* cDNA from control and MO-induced Hpa2 LOF. Arrows point towards splice variants. **(I)** Edema scoring in MO-induced Hpa2-LOF larvae. **(J)** Quantification of vascular integrity in MO-induced Hpa2-LOF larvae at 96 hpf with increasing MO concentrations. (**K**) Quantification of vascular permeability in MO-induced Hpa2-LOF larvae with human *hpse2* mRNA co-injection. **(L)** Transmission election microscopy of the glomerular filtration barrier in Hpa2-LOF larvae at 96 hpf. Asterisks indicate swollen ECs, hashtags indicate basement membrane (BM) infolding, FP = podocyte foot processes (arrowheads). (**M**) Hpa2 LOF in *Tg(fli1:EGFP)* zebrafish induces occasional abnormal sprouting (arrowheads) of intersegmental vessels (ISV) in zebrafish tail vasculature at 72 hpf. (**N**) Proportion of fish with abnormal sprouting in ISV. (**O**) Diameter along ISV in Hpa2-LOF scenarios. Data points represent individual analyzed fish. All quantitative data were generated from at least two independent inductions of Hpa2 LOF. Multiple comparisons by one-way ANOVA and Tukey multiple comparison tests. Pairwise comparisons by paired t-test. ****p ≤ 0.0001; ***p ≤0.001, ** p ≤0.01; * p ≤0.05; n.s. = not significant. Bar height shows the mean ± SD.

### Hpa2 LOF causes morphological changes in glomerular ECs and changes blood-vessel diameter

The kidney is the primary filtrating organ in vertebrates and has a very complex and delicate vascular architecture. HS plays an important role in maintaining the integrity of the glomerular filtration barrier (GFB), and frequently protein is lost via leakage through the vasculature of glomeruli^31^. Hence, we hypothesized that Hpa2-LOF may disrupt GFB architecture, thereby exacerbating the vascular-leakage phenotype in the kidney. To test this, we performed an in-depth analysis of the GFB by transmission electron microscopy. The GFB of Hpa2-LOF larvae showed pathological abnormalities characterized by swollen ECs, which lost their fenestrated morphology and basement-membrane infolding, whereas podocyte morphology showed no abnormalities (Figure 2L, Figure S3D). Hpa2-LOF larvae also exhibited reduced staining of the glomerular capillary space and a slightly increased staining of the urinary space between podocyte foot processes (Figure 2L, Figure S3D), suggesting leakage of plasma from capillaries into the urinary space. We noticed no changes in tubular morphology (data not shown). Together, these findings argue for a loss of protein from the glomeruli.

To investigate whether the impact of Hpa2 LOF on the vasculature is not limited to the kidney, we screened vessel morphology in EC-labeling *Tg(fli1:EGFP)* zebrafish by analyzing intersegmental vessels (ISV) of the tail vasculature. Hpa2 LOF induced a mild phenotype in the tail vasculature characterized by formation of occasional sprouts in 33% of investigated larvae (Figure 2M, N). Additionally, the vessel diameter of ISV was 9.22 µm in Hpa2-LOF larvae compared to 8.34 µm in control-injected animals (Figure 2O). Concordantly, vessel diameter soared to 9.17 µm in MO-induced Hpa2-LOF larvae compared to 8.43 µm in control-injected animals (Figure 2O). Together, these data provide evidence for an Hpa2-LOF-induced vascular phenotype characterized by vascular hyperpermeability, changes in EC and vessel morphology, and a mild pro-angiogenic phenotype.

### Recombinant human Hpa2 (hHpa2) rescues the Hpa2-LOF phenotype in zebrafish

In our next set of experiments, we aimed to better understand the effect of Hpa2 on endothelial HS. We predicted the three-dimensional structure of human Hpa2 (hHpa2) using AlphaFold2 software and calculated the electrostatic surface potential, which revealed large patches of positive electrostatic potential on the protein surface (Figure 3A). We then modeled insertion of a heparin tetrasacchride ligand into the predicted three-dimensional structure of Hpa2, using the ClusPro server, which has been shown to reliably predict heparin-binding sites based on clustering of docked structures^32^. The modeling located the cluster with the lowest energy (675 members; lowest energy score = −1069.3) in the positively charged electrostatic area of hHpa2 (Figure 3A, B). Major contributors to the interaction are predicted to be the basic amino acids Arg466, Lys477, Arg506, Arg508, Lys509, Lys510, and Arg567 (Figure 3B). All the residues are conserved between human and zebrafish (Figure 1B).

**Figure 3:**
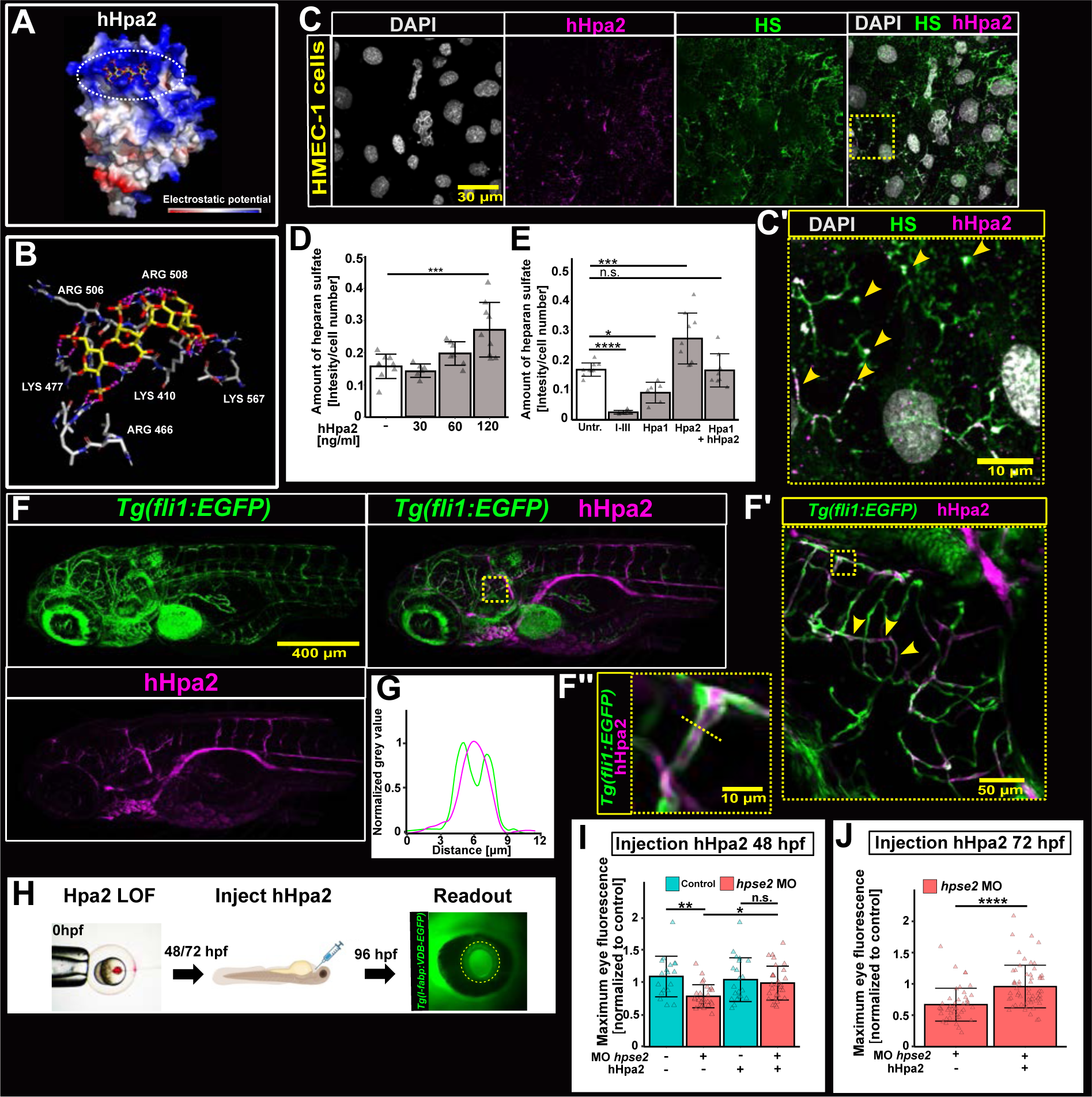
Recombinant human Hpa2 (hHpa2) increases endothelial heparan sulfate (HS) and alleviates the Hpa2 LOF phenotype in zebrafish. **(A)** Electrostatic surface potential of hHpa2 (amino acids 60-592) with predicted heparin tetrasaccharide ligand (structure within white circle) binding. Blue indicates positive surface potential, red negative. **(B)** Putative ionic interactions (magenta) of basic hHpa2 residues (white carbon scaffold) and heparin (yellow carbon scaffold). Oxygen in red, nitrogen in blue, sulfur in bronze. **(C)** Immunofluorescence of added hHpa2 (magenta) and HS (green) in human microvascular ECs (HMEC-1). C’: Arrowheads indicate overlapping hHpa2 and HS signals. **(D,E)** Quantification of HS signal intensity 4 h after the addition of hHpa2 (D) or of (E) HS-degrading enzymes heparinases I-III (I-III) or heparanase 1 (Hpa1), or Hpa1 together with hHpa2, to HMEC-1 cells. Datapoints refer to individual slides collected from three independent stainings. Bar height shows the mean ± SD. **(F, F’, F”)** Whole-mount immunofluorescence staining for hHpa2 in *Tg(fli1:EGFP)* larvae 12 h after injection of hHpa2. Arrowheads in ROI _indicate_ overlapping antibody signal for V5-tagged hHpa2 and EGFP (F’, F’’). Yellow dotted line in F” shows region of vessel lumen analyzed in G. **(G)** Intensity plot for hHpa2 detected by a V5 antibody and EGFP signal across vessel lumen indicated in F’’. Intensity was normalized to the maximum intensity in each channel. **(H)** Treatment schedule for hHpa2 injections (86 pg) into Hpa2-LOF *Tg(l-fabp:DBP-EGFP)* larvae at 48 hpf or 72 hpf. (**I, J)** Quantification of vascular permeability after injection of hHpa2 at 48 hpf (I) and 72 hpf (J). Datapoints show individual larvae, bar height shows the mean +SD per group. Multiple comparisons by one-way ANOVA and Tukey multiple comparison tests. Pairwise comparisons by paired t-test. ****p ≤ 0.0001; ***p ≤0.001, ** p ≤0.01; * p ≤0.05; n.s. = not significant. Bar height shows the mean ± SD.

We developed a stable mammalian expression system to generate recombinant hHpa2 with a C-terminal V5-tag. hHpa2 was purified using a heparin-based chromatographic procedure (Figure S4). Using this system, we then validated the functionality of purified hHpa2 in human microvascular EC (HMEC-1) cultures by colocalizing the recombinant protein with HS on the EC surface (Pearson’s coefficient = 0.373; Figure 3C). Further, the addition of hHpa2 increased the amount of EC HS in a dose-dependent manner, resulting in an increase of 72% in HS after 4 hours of stimulation with 120 ng hHpa2 (Figure 4D). The addition of bacterial heparinases I-III and mammalian Hpa1 resulted in decreased HS on the EC surface (Figure 3E). We confirmed that hHpa2 inhibits the HS-degrading action of Hpa1 (Figure 4E). From these experiments we conclude that the generated recombinant hHpa2 is fully functional and that it has the capacity to protect and maintain HS on the EC surface. We next used hHpa2 to test for a cross-species rescue by injecting the human protein into Hpa2-LOF zebrafish larvae. First, we injected the protein into the cardinal vein of *Tg(fli1:EGFP)* larvae at 84 hpf. Twelve hours after injection, hHpa2 was still present in the larvae and localized in the lumen of blood vessels, which we detected with a V5 antibody to discriminate hHpa2 from endogenous Hpa2 (Pearson’s coefficient =0.308; Figure 3F, G). Next, we injected hHpa2 into Hpa2-LOF *Tg(l-fabp:DBP-EGFP)* larvae to test its capacity to rescue the vascular-leakage phenotype. hHpa2 was injected at either 48 hpf, i.e., before the endogenous expression of hepatic *hpse2*, or at 72 hpf, after endogenous Hpa2 naturally circulates in the vasculature (Figure 3H). hHpa2 alleviated the Hpa2-LOF phenotype, increasing the MEF to 96% compared to 76% in control-injected larvae (Figure 3I). After the injection of hHpa2 into Hpa2-LOF larvae at 72 hpf, the MEF soared from to 96% compared to 67% in controls (Figure 3H). Together, these results lead us to conclude that hHpa2 increases vascular HS and that the role of Hpa2 is functionally conserved in the vasculature of zebrafish and humans.

**Figure 4:**
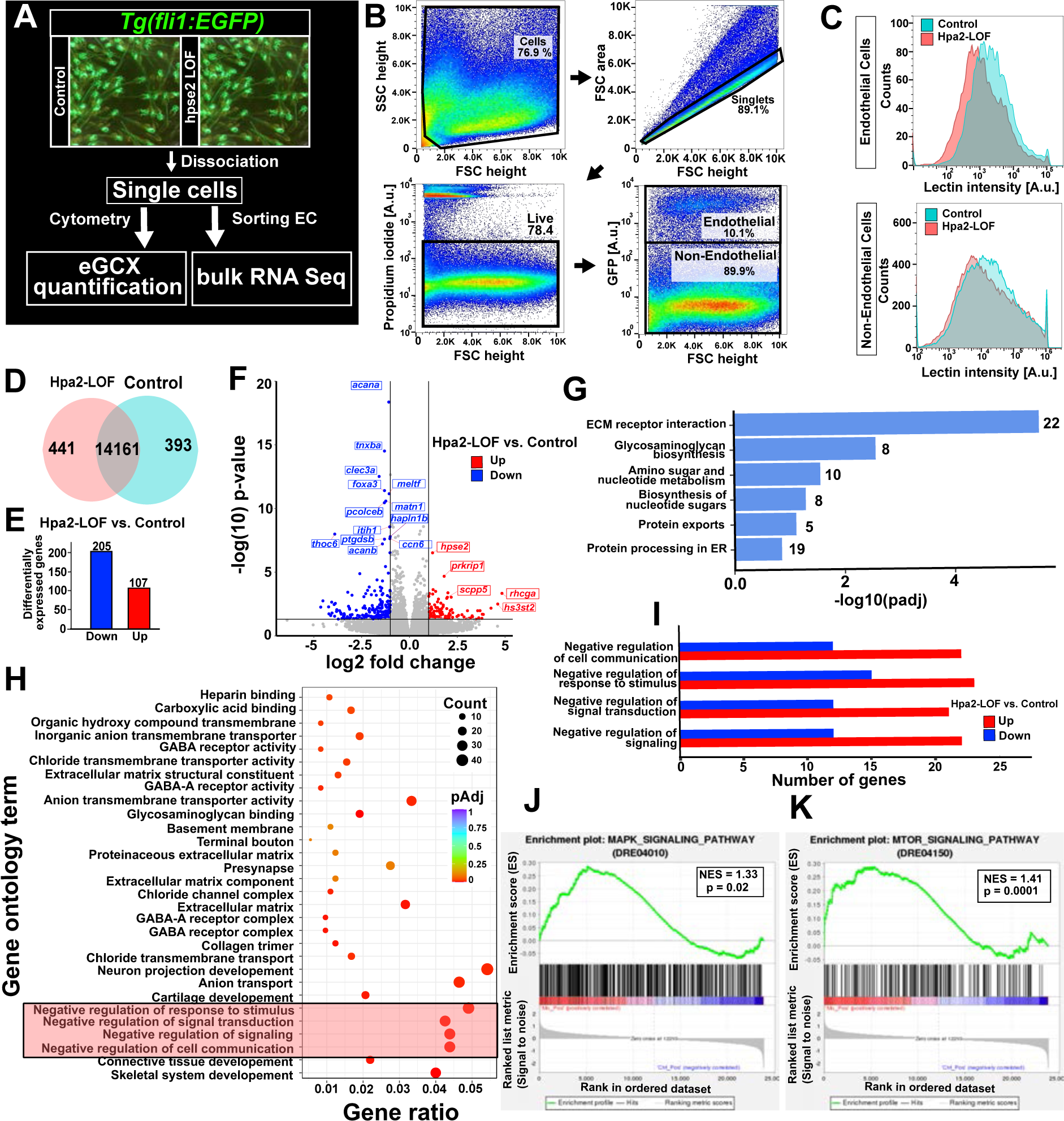
Hpa2 LOF reduces endothelial glycocalyx (eGCX) quantity and dysregulates EC signaling in zebrafish larvae. **(A)** Workflow for EC analysis in Hpa2-LOF larvae with a *Tg(fli1:EGFP)* background. Hpa2-LOF fish and control fish were pooled and dissociated into single cells at 96 hpf. Cells were either stained and subjected to quantitative analysis using cytometry, or were sorted to enrich GFP**^+^** ECs. RNA of ECs was isolated and subjected to bulk RNA sequencing. **(B)** Gating strategy for sorting and analyzing ECs. **(C)** Quantitative analysis of GCX stained by wheat germ agglutinin in GFP**^+^** cells (upper) and non-GFP**^+^** cells (lower) comparing Hpa2-LOF and control larvae at 96 hpf. **(D)** Venn diagram of genes expressed in Hpa2-LOF (red) and control (blue) larvae. (**E**) Number of genes differentially expressed (red: upregulated; blue: downregulated) between Hpa2-LOF and control ECs. **(F)** Volcano plot of differentially expressed genes. Genes with p value ≤ 0.05 and fold change ≥ 2 are colored. **(G)** Kyoto Encyclopedia of Genes and Genomes (KEGG) analysis of differentially regulated pathways. The numbers to the right of the horizontal bars indicate the numbers of differentially regulated genes in the corresponding pathway. **(H)** Gene Ontology analysis of dysregulated biological processes. Gene ratio expresses the number of differentially expressed genes in a pathway relative to all genes grouped to that pathway. **(I)** Number of genes that are up- (red) or down-regulated (blue) in the corresponding biological process. **(J,K)** Gene set enrichment analysis of MAPK (mitogen-activated protein kinase) signaling pathway (J) and mTOR (mammalian target of rapamycin) signaling pathway (K). NES = Normalized enrichment score.

### Hpa2 LOF in zebrafish causes changes in the eGCX and alters the response of ECs to signaling molecules

We next asked how Hpa2 LOF affects EC physiology on a molecular level *in vivo*. We addressed this by analyzing ECs from *Tg(fli1:EGFP)* larvae. Hpa2-LOF or control-injected larvae were pooled and then dissociated into single cells, after which GFP^+^ ECs were either subjected to cytometry analysis for investigation of cell-surface eGCX structures, or sorted to enrich GFP^+^ ECs for bulk RNA sequencing (Figure 4A). Dissociated cells were examined for cellular viability (80% alive) before gating for GFP^+^ EC (10% of live cells, Figure 4B). We stained ECs with wheat germ agglutinin (WGA) to quantify cell-surface N-acetylglucosamine and sialic acid. We detected a 15.5% decrease in lectin binding in Hpa2-LOF ECs compared to ECs derived from control-injected larvae (Figure 4C, Figure S5A). WGA binding to GFP-negative cells showed no significant difference between Hpa2-LOF and control cells (Figure 4C, Figure S5B). Transcriptomes of enriched ECs derived from MO-induced Hpa2-LOF and control larvae were analyzed for differentially expressed genes. Numerous genes were exclusively expressed in either MO-derived or control-derived ECs (Figure 4D). Compared to control-injected larvae, Hpa2 LOF caused downregulation of 295 genes and upregulation of 107 genes (Figure 4E, F). The most significantly upregulated gene in enriched ECs of Hpa2-LOF larvae was *hpse2* (Figure 4F). Pathway analysis by KEGG identified dysregulated expression of genes involved in extracellular-matrix-receptor interaction including genes that encode collagens, laminins, and thrombospondins (Figure 4G, Figure S5C), and genes involved in GAG biosynthesis (chondroitin sulfate and HS) and amino sugar and nucleotide metabolism (Figure 4G, Figure S5D). Gene Ontology analysis of biological processes revealed changes in genes involved in negative regulation of response to stimuli; of signal transduction; and of cell-cell communication (Figure 4H). Gene clusters of these biological processes were primarily upregulated (Figure 4I). ECs rely heavily on finely regulated mitogen-activated protein kinase (MAPK) signaling and phosphatidylinositol 3-kinase (PI3K)-Akt signaling to maintain a homeostatic state^1^. Because genes involved in overall signal transduction were strongly dysregulated in ECs of Hpa2-LOF larvae, we wanted to know whether these two signaling pathways were affected. Gene set enrichment analysis revealed enrichment of genes involved in MAPK signaling and the Akt-mammalian target of rapamycin (mTOR) signaling pathway (Figure 4J, K, Figure S5E,F), suggesting dysregulated activation of these pathways in Hpa2-LOF-derived ECs. MAPK and Akt signaling in ECs are activated primarily by members of the receptor tyrosine kinase (RTK) family, including VEGFA –VEGF receptor 2 (VEGFR2) signaling and FGF–FGF receptor (FGFR) signaling. Both ligand families contain members with HS-binding domains and are regulated by endothelial HS^33^.

### hHpa2 blunts EC signal response towards FGF2 and VEGFA by reducing ligand binding to the EC surface

We previously observed strong binding affinity of Hpa2 towards HS (Figure 3 A-E, Figure S4), and thus hypothesized that Hpa2 can modulate HS-dependent signaling pathways. We first tested this *in vitro*, using HMEC-1 cells, which were shown above to form stable amounts of HS under static conditions (Figure 3C). We chose to use FGF2 and VEGFA_165_, as they are two of the most important growth factors for EC physiology, each contains HS-binding domains^1^. Both growth factors signal through RTK, which activates the MAPK and PI3K pathways for EC migration, proliferation, and survival^34,35^. First, we tested binding of these growth factors on the EC surface in the presence and absence of hHpa2. We confirmed the known dependence of FGF2 and VEGFA_165_ binding on the presence of HS^36,37^ by showing decreased binding of these factors when cells were pretreated with HS-degrading bacterial heparinases (Figure 5A, B). Specifically, compared to non-pretreated cells, FGF2 binding was reduced by 53% and VEGFA binding by 22% when cells were pretreated with heparinases I-III (Figure 5A, B). Treating the cells with hHpa2 and growth factors reduced growth-factor binding in a dose-dependent manner; the application of 24 ng hHpa2 compared to cells not treated with hHpa2 decreased FGF2 binding by 43% (Figure 5A) and decreased VEGFA binding by 23% (Figure 5B). Next, we tested the intensity of signal transduction elicited by these factors in the presence and absence of hHpa2. hHpa2 reduced FGF2-induced extracellular receptor kinase 1/2 (ERK1/2) phosphorylation in a dose-dependent manner, reaching a maximum decrease of 41% in phosphorylation (Figure 5D). Modeling a function for the hHpa2-mediated decrease in ERK phosphorylation confirmed the correlation and revealed a logarithmic relationship (r^2^= 0.55, Figure 5E). Consistently, treating the cells with at least 90 ng hHpa2 reduced VEGFA_165_-induced phosphorylation of VEGFR2 at Tyr1175 by 23% (Figure 5F, G). We did not observe a statistically significant decrease in receptor phosphorylation using less than 60 ng hHpa2 (Figure 5F, G). To further confirm these findings, we performed immunofluorescence staining using a phospho-tyrosine antibody to quantify cell activation after growth-factor and Hpa2 application. hHpa2 decreased the intensity for both FGF2- (Figure 5H, I) and VEGFA_165_- (Figure 5J, K) induced p-tyrosine activation by 20%. Accordingly, phosphorylation of VEGFR2 at Tyr1175 by VEGFA_165_ was reduced by 20% in the presence of hHpa2 (Figure 5J, L). Together, these data provide evidence that hHpa2 competes with growth factors for HS binding, thereby reducing the signaling input elicited by these factors in ECs.

**Figure 5:**
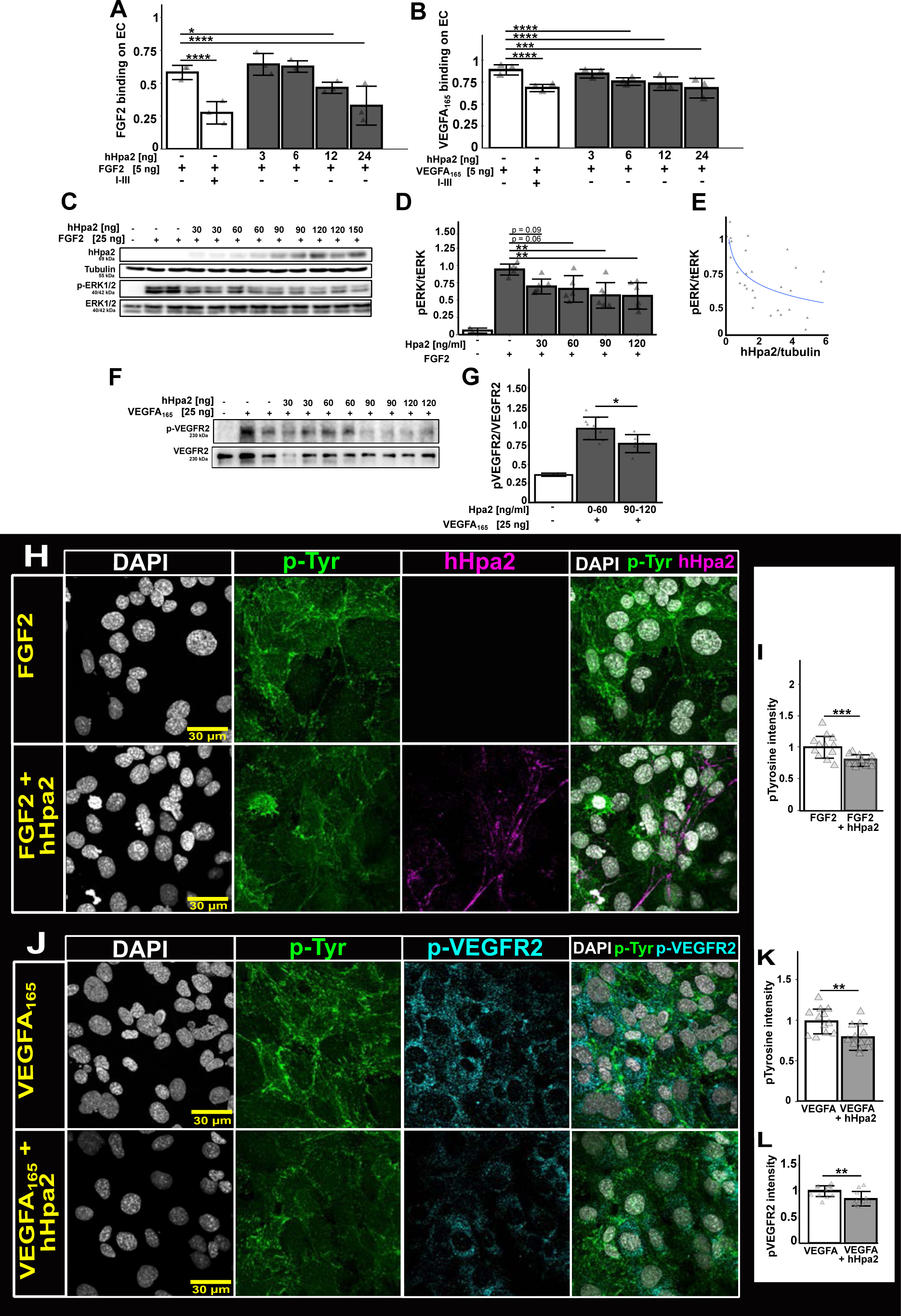
Recombinant human Hpa2 (hHpa2) reduces binding and signal induction of HS-binding growth factors in HMEC-1. **(A, B)** Recombinant growth-factor binding of FGF2 (fibroblast growth factor 2) (A) and VEGFA_165_ (vascular endothelial growth factor A_165_) (B) to HMEC-1 cells in the presence and absence of hHpa2. Data are normalized to binding of recombinant growth factor without hHpa2 and corrected for signal from endogenously synthesized FGF2 or VEGFA_165_. Datapoints show three independent biological experiments, each in triplicate. **(C)** Representative western blot of cell lysates stimulated with FGF2 (25 ng) in the presence and absence of hHpa2. **(D)** Quantification of phospho-ERK1/2 (pERK1/2) activation by FGF2 after 5 min. **(E)** Correlation of pERK/tERK and hHpa2 intensities after FGF2 stimulation. **(F)** Representative western blot of cell lysates stimulated with VEGFA_165_ (25 ng) in the presence and absence of hHpa2 after 10 min. **(D)** Quantification of phospho-VEGFR2 (p-VEGFR2) activation by VEGFA_165_. **(H, J)** Phospho-Tyrosine (p-Tyr) activation in HMEC-1 cells after stimulation with 25 ng FGF2 (H) or VEGFA_165_ (J) without (upper) and with hHpa2 (lower) after 5 min (FGF2) and 10 min (VEGFA_165_). **(I, K, L)** Quantified intensities of p-Tyr (I, K) and p-VEGFR2 (L) activation. Datapoints refer to individual frames collected from three independent stainings. Bar height shows the mean ± SD. Multiple comparisons by one-way ANOVA and Tukey multiple comparison tests. Pairwise comparisons by paired t-test. ****p ≤ 0.0001; ***p ≤0.001, ** p ≤0.01; * p ≤0.05; n.s. = not significant. Bar height shows the mean ± SD.

### Inhibition of RTK signaling in zebrafish mitigates the Hpa2-LOF phenotype

Finally, we tested the principle of Hpa2 as a modulator of HS-dependent signaling for ECs *in vivo*. We treated Hpa2-LOF larvae with different drugs known to interfere with both FGF and VEGF signaling. Competitive inhibition of VEGFR2/FGFR by brivanib alleviated the vascular hyperpermeability phenotype of Hpa2-LOF larvae by 52%, shifting the MEF from 77% in control larvae to 90% in Hpa2-LOF larvae (Figure 6A,B). SU5402, another RTK inhibitor but more specific for FGFR1, alleviated the vascular hyperpermeability in Hpa2-LOF larvae by 58%, with an increase in MEF from 74% in control larvae to 90% in Hpa2-LOF larvae (Figure 6C, D). Next, we directly inhibited MAPK signaling by U0126, an inhibitor specific for mitogen-activated protein kinase kinase, the upstream kinase of ERK. U0126 alleviated the permeability phenotype by 35%, with the MEF rising from 61% in control larvae to 75% in Hpa2-LOF larvae (Figure 6E, F). Notably, application of U0126 showed tendencies to reduce the MEF in control larvae (Figure 6E). Lastly, we tested LY294002, an inhibitor specific for the PI3K/Akt pathway. LY294002 did not change the phenotype of Hpa2-LOF larvae (Figure 6G, H), suggesting that the permeability phenotype is caused primarily by the MAPK pathway. Together, these data demonstrate that the hyperpermeability phenotype of Hpa2-LOF in zebrafish larvae is caused by overactivation of FGF/VEGFA-MAPK signaling. This in turn leads us to establish the model of Hpa2 as a circulating molecule that regulates HS-dependent processes on the EC surface, including HS degradation and HS-dependent RTK signaling (Figure 7).

**Figure 6:**
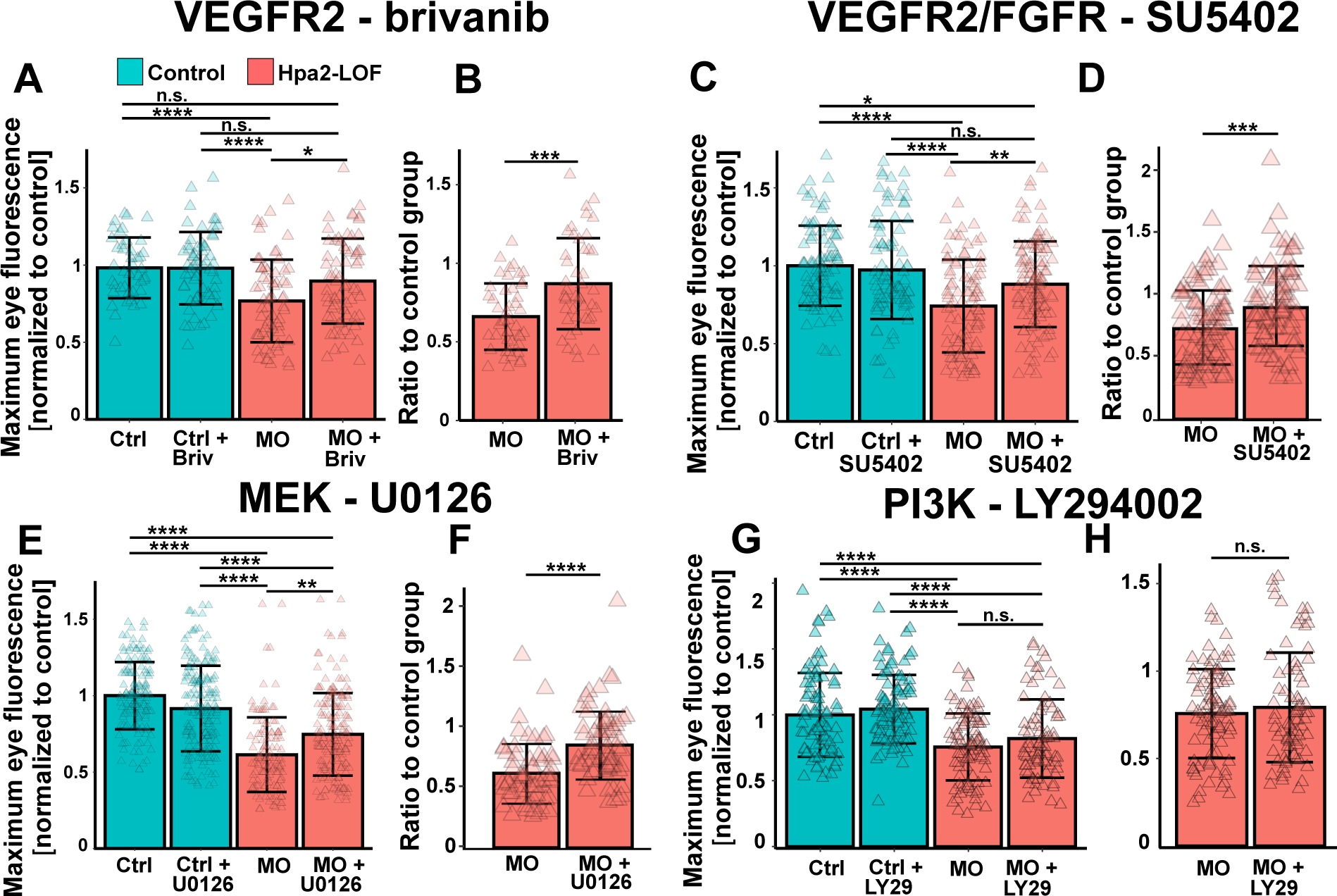
Inhibition of growth-factor-response pathways alleviates hyperpermeability phenotype in Hpa2-LOF larvae. **(A)** Maximum eye fluorescence (MEF) of brinvanib-treated (10 nM) Hpa2-LOF larvae (0.58 pmol *hpse2* MO) at 96 hpf**. (B)** Ratio of MEF after brivanib treatment compared to control counterparts. **(C)** MEF of SU5402-treated (250 nM) Hpa2-LOF larvae (0.58 pmol *hpse2* MO) at 96 hpf**. (D)** Ratio of MEF after SU5402 treatment compared to control counterparts. **(E)** MEF of U0126-treated (100 nM) Hpa2*-*LOF larvae (0.58 pmol *hpse2* MO) at 96 hpf**. (F)** Ratio of MEF after U0126 treatment compared to MEF of control counterparts. **(G)** MEF of LY294002-treated (1000 nM) Hpa2*-*LOF larvae (0.58 pmol *hpse2* MO) at 96 hpf **(H)** Ratio of MEF after LY294002 treatment compared to MEF of control counterparts. Multiple comparisons by one-way ANOVA and Tukey multiple comparison tests. Pairwise comparisons by paired t-test. ****p ≤ 0.0001; ***p ≤0.001, ** p ≤0.01; * p ≤0.05; n.s. = not significant. Bar height shows the mean ± SD.

**Figure 7:**
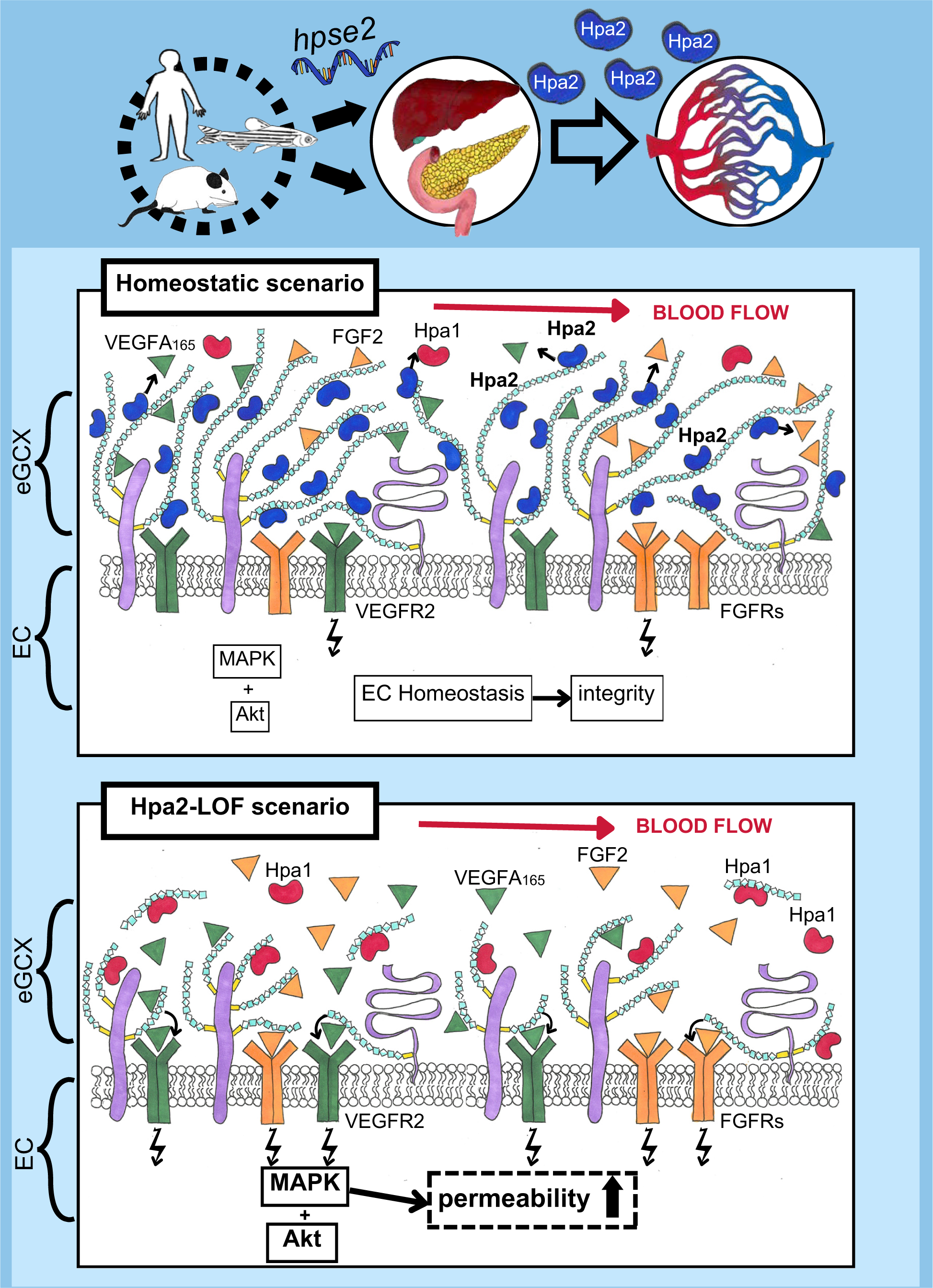
Model: Hpa2 as a circulating regulatory factor for the vascular system. The *hpse2* gene is expressed primarily by secretory organs. Hpa2 protein is released into the vasculature, where it binds HS on the EC surface. Hpa2 competes with HS-binding proteins including Hpa1, FGF2, and VEGFA_165_ for HS-binding sites. This reduces the amount of growth factors locally available on the EC surface and translates into decreased receptor activation by these ligands. Loss of Hpa2 increases Hpa1-induced HS shedding and promotes binding of HS-dependent growth factors FGF2 and VEGFA_165_, which subsequently increases vascular permeability.

## Discussion

In this study we characterized the role of Hpa2 for in cardiovascular system using zebrafish as our primary research model. We describe Hpa2 as a novel regulator of EC function that controls HS-dependent processes. Functional loss of Hpa2 caused EC dysfunction with increased vascular permeability and vessel dilatation, and aberrant EC morphology. Lack of Hpa2 led to overactivation of MAPK signaling in ECs. Hpa2-LOF induced rarefaction of the eGCX and a change in receptor-mediated signaling. Our findings indicate that Hpa2 is a key component of a homeostatic system protecting the endothelium and thus may play a role in EC disease and have substantial potential as a therapeutic target.

Over the last decade, studies have begun to elucidate the physiological function of Hpa2, describing roles primarily in cancer biology and development of the peripheral nervous system^38^. Only recently have studies shown Hpa2 to play a protective role in the vascular system. However, these latter studies were based on the exogenous application of Hpa2^23^ or use of overexpression systems^22^ and were performed under disease conditions. Here, we characterize for the first time the role of endogenous Hpa2 in the systemic vasculature and find that Hpa2 is a homeostatic regulator of HS-dependent processes and EC physiology.

### Hpa2 as a regulator of vascular permeability

Hpa2-LOF in zebrafish larvae caused increased vascular permeability, vessel dilatation, and altered EC morphology (Figure 2J, L, M), providing evidence that ECs are significantly affected by a functional loss of Hpa2. Hpa2 LOF decreased eGCX quantity *in vivo* (Figure 4C), and hHpa2 increased HS abundance *in vitro* (Figure 5D). The physiological role of the eGCX in maintenance of vascular integrity, particularly in permeable vascular beds such as the glomeruli of kidneys, is well characterized^31^ and is reinforced by the findings in this study. Hpa2 LOF caused a significant endothelial phenotype in the GFB accompanied by protein leakage into the urinary space (Figure 2L), underlining the importance of Hpa2 and the eGCX for glomerular function. Interestingly, *hpse2^-/-^* mice showed mild albuminuria and markers for kidney dysfunction during their short lifetime^26^. Because this mouse study focused primarily on the bladder phenotype, it remains to be determined whether the kidney phenotype observed in mice arises by vesicoureteral reflux, which damages the kidney from the urinary space, or whether the phenotype arises from the vascular side, as we observed here in zebrafish. We could not find any *hpse2* expression in kidney-associated cell types, implying that circulating, liver-derived Hpa2 provides the nephroprotective effects. This is supported by previous studies that applied exogenous Hpa2 to alleviate pathological kidney phenotypes^22,23^ and by our finding that hHpa2 alleviates the Hpa2-LOF phenotype in zebrafish (Figure 3I, J). These results do not rule out the possibility that protein leakage also occurs in other regions of the body. Our observations of dilated vessel diameter and occasional sprout formation in Hpa2-LOF larvae (Figure 2M, N), as well as *hpse2* expression in endocrine organs (Figure 1D-E, Figure S1), argue for a more systemic role of Hpa2. However, due to the fragility and higher permeability of the fenestrated endothelium in the glomeruli, we reason that loss of Hpa2 has a strong impact on the kidney vasculature and that a large proportion of protein is lost from this region.

### Regulation of Hpa2 expression

We consistently observed *hpse2* expression in central endocrine organs such as the liver and pancreas across the examined vertebrates (Figure 1D-E, Figure S1). Hpa2 therefore has the characteristics of an endocrine molecule acting on the systemic vasculature, which immediately raises questions about its regulation. For example, a previous study classified *hpse2* as a stress-response gene that is transcriptionally induced by endoplasmic-reticulum stress and hypoxia^39^. The authors identified activating transcription factor 3 as a major regulator of *hpse2* expression in cancer-cell lines. Activating transcription factor 3 is activated by a range of stress stimuli including metabolic and inflammatory signals and is an important regulator of endocrine organs such as liver and pancreas^40^. Thus, *hpse2* may be a direct responsive gene of activating transcription factor 3 in the liver and pancreas, induced under systemic stress conditions. We noted *hpse2* upregulation in ECs of Hpa2-LOF larvae (Figure 4F), implying the adaptability of ECs in expressing their own *hpse2* if it is not supplied by the circulation. Further studies of *hpse2* expression regulation under physiological and pathophysiological stress conditions are needed to fully understand the systemic regulatory function of Hpa2 in an organism.

### Hpa2 has a conserved role in neural development

Hpa2 deficiency in humans is linked to urofacial syndrome, an autosomal recessive disorder characterized by urological and facial abnormalities^41,42^. The phenotype of Hpa2-deficient mice largely resembles that of Hpa2-deficient humans, showing abnormal bladder innervation and growth retardation, ultimately resulting in death within the first weeks after birth^25,26,43^. A role for Hpa2 in neural development is also indicated by a study showing neural expression of *hpse2* in *Xenopus* development accompanied by overactivated MAPK signaling^29^. This is consistent with our findings of early *hpse2* expression in dorsal root ganglia along the spinal cord (Figure S2A, B) and overactivated MAPK signaling in Hpa2-LOF larvae (Figure 4J). Our observation of decreased survival in F0 CRISPR-Cas9 induced Hpa2-LOF larvae (Figure S3B) and our inability to breed homozygous *hpse2* mutants suggest that a total loss of Hpa2 causes embryonic lethality in zebrafish. Together, these studies demonstrate that Hpa2 is necessary for proper development of the peripheral nervous system across vertebrates and that its complete absence is incompatible with life.

Compared to embryos injected with control MO, embryos with MO-induced Hpa2-LOF exhibited no significant decrease in survival or pericardial edema formation (Figure S3B), which were the only phenotypes that differed between MO-induced Hpa2-LOF and CRISPR-Cas9 LOF embryos. It is likely that the *hpse2* MO does not cause a total loss of functional Hpa2 protein, and that once hepatic expression of *hpse2* starts, i.e., between 60 and 72 hpf, the MO is already too diluted to cause sufficient Hpa2-LOF to induce edema formation. Generally, the CRISPR-Cas9-induced Hpa2 LOF approach showed a slightly stronger phenotype compared to the MO approach (Figure 2). However, further elucidation of the role of Hpa2 in different organs and at different developmental stages will require tissue-specific controllable models to knock out and overexpress Hpa2.

Recently, a pan-inducible knockout of Hpa2 was generated in mice that caused a pancreatic phenotype characterized by decreased organ weight, increased pancreatic inflammation, and morphological alterations^44^. No urinary-tract phenotype was reported in these mice, which argues that in adulthood Hpa2 plays a negligible role in the nervous system but is important for the function of other organs including the pancreas and kidney. Concordantly, we detected *hpse2* expression in the murine pancreas (Figure S1A, B). Future studies assessing the pancreatic and glomerular vasculature in these animals with a focus on the eGCX and the inflammatory state of the endothelium would help to further understand the role of Hpa2 in organ-specific vascular homeostasis. It would be very interesting to explore whether Hpa2-deficient patients and mice exhibit eGCX impairment and increased levels of eGCX degradation products in their blood, and show increased susceptibility to environmental stress stimuli.

### Hpa2 as a regulator of HS-dependent EC signaling

Endothelial quiescence and normalcy is defined as a non-proliferative, non-angiogenic, and resilient state that requires a defined signal signature^1^. Signaling input must be tightly regulated so that ECs are maintained in a non-proliferative and anti-angiogenic state but at the same time do not undergo apoptosis and cell death due to signal starvation^45,46^. ECs use pericellular HS to regulate the number of spatiotemporally available ligands for their receptors on the cell surface^47^. Signaling of these factors is significantly impaired in the absence of HS^48^. Here we show that Hpa2 reduces binding of the HS-binding growth factors VEGFA_165_ and FGF2 and consequently reduces the intensity of signal transduction elicited by these factors (Figure 5C-G). We and others previously showed that Hpa2 binds HS with strong affinity^19^. The ability of Hpa2 to displace VEGFA_165_ and FGF2 from the EC surface might be a systemic mechanism to regulate HS-dependent signaling in ECs. The concept of competitive growth-factor release from ECM-bound reservoirs is long known and was recently demonstrated in a study showing the on-demand release of VEGFA_165_ from the ECM in zebrafish spinal-cord vascularization, with the release of VEGFA_165_ inducing angiogenic sprouting towards the VEGFA_165_ gradient^49^. We did not observe prominent accumulation of Hpa2 in the interstitium. Rather, Hpa2 showed marked expression in hepatic tissue and vascular localization among all examined species (Figure 1, Figure S1). Thus, competition for HS binding likely takes place at the ligand’s destination, i.e., the EC surface.

Transient activation of ECs by VEGFA results in vascular permeability and vessel relaxation^50^ while chronic overstimulation induces vascular malformations^51^. Additionally, levels of VEGFA are finely controlled in the GFB and disturbance of the system, including up- or downregulation of VEGFA, cause a severe endothelial phenotype in glomeruli^52^. Similar effector functions were observed in our Hpa2-LOF model, including increased vascular permeability and vessel dilatation, and changes in EC morphology (Figure 2K, L, O). Maintenance of vascular integrity and endothelial identity in the quiescent state are also regulated by the FGF system^53,34^. Evidence that Hpa2 acts as a regulator for the VEGFA and FGF signaling pathways on the EC surface in zebrafish is provided by several of our observations in Hpa2-LOF larvae: 1) MAPK signaling genes are enriched (Figure 4L); 2) genes involved in negative regulation of signaling are enriched (Figure 4J, Figure S5E); 3) pharmacological inhibition of VEGFR2/FGFR/MAPK signaling alleviated vascular permeability (Figure 6A-F); and 4) FGF2/VEGFA_165_ displacement was mediated by Hpa2 (Figure 5A, B). Consistent with these observations, Hpa1-mediated HS degradation induces the release of FGF2 from the ECM and promotes angiogenesis in ECs^54^. Inhibition of the Akt/mTOR pathway by LY294002 did not alleviate the vascular-leakage phenotype induced by Hpa2 LOF (Figure 6G, H), although gene-expression profiles indicated overactivation of this pathway (Figure 4K, Figure S5F). In ECs, PI3K/Akt signaling is primarily responsible for cell survival and migration, whereas ERK1/2 signaling is primarily responsible for control of vascular integrity^55^. It is therefore reasonable that inhibition of PI3K does not alleviate vascular leakage, as we observed (Figure 6G, H).

Our study has revealed important new information on the specificity with which Hpa2 impacts the binding of HS-binding growth factors. For example, our cell binding assays show that the impact of Hpa2 on HS binding of FGF2 is stronger than its impact on HS binding of VEGFA_165_, as bacterial heparinases I-III impaired FGF2 binding on ECs more strongly than they impaired VEGFA_165_ binding (Figure 5A, B). In addition, we can predict that Hpa2 has a particularly strong impact on signaling induced by HS-binding proteins that bind the same HS domains that Hpa2 binds. However, much remains unknown regarding the specificity with which Hpa2 competes with other HS-binding binding proteins including growth factors. To more thoroughly understand the effects of Hpa2 on HS-dependent processes, assays characterizing the binding of different HS/GAG species with Hpa2, and computational modeling, are required. In addition, site-directed mutagenesis studies are needed to investigate whether Hpa2 harbors additional HS-independent functions.

### A novel strategy to protect and preserve the eGCX using Hpa2

The eGCX plays major roles in vascular homeostasis and vascular health. Restoration of the eGCX has been associated with symptom alleviation in microvascular diseases including chronic kidney disease^56,57^, diabetic microangiopathy^58^, and sepsis^59^. Thus, therapeutic strategies targeting the eGCX would have a significant impact on human health. Hpa1 represents a particularly attractive therapeutic target for such a strategy. A direct effect of Hpa1 on microvascular disease progression was demonstrated in a study showing that *hpse^-/-^* mice show reduced susceptibility to development of diabetes-induced albuminuria^16^. More recently, it was demonstrated that Hpa1 inhibition protects from hyperglycemia-induced glomerular eGCX degradation in *db/db* mice, a model of type 2 diabetes, indicating the potential of Hpa1 inhibition as a therapeutic strategy for microvascular disease^58^. Inhibitors of Hpa1 have been studied primarily to treat neoplasias^60^. However, to date Hpa1 inhibitors have failed in clinical trials, likely because they are structurally related to heparin, which is known for its adverse pleiotropic effects, including anticoagulatory activity, that can easily lead to poor outcomes in critically ill tumor patients^61^. Here we show, for the first time, that an intact eGCX and a homeostatic vascular state are dependent on Hpa2, indicating that Hpa2 may provide a novel and effective treatment strategy for microvascular disease and cancer. Importantly, because Hpa2 does not interact directly with Hpa1, it avoids the adverse effects of heparin mimetics and provides beneficial pleotropic effects for the vasculature by reducing the amount of HS-dependent signal transduction in ECs (Figures 4, 5). Inhibiting angiogenesis and endothelial activation has been widely practiced in modern medicine by inhibition of the HS-dependent VEGFR2 pathway in retinal neovascularization and tumor vascularization^62^. This decreased signal transduction would also be advantageous in cancer treatment, as it would be expected to reduce EC proliferation and migration. The first evidence of exogenous Hpa2 application for the treatment of vascular disease was published recently^22,23^. Here, we expand the potential therapeutic spectrum of Hpa2 by providing evidence that the effects of Hpa2 do not rely solely on the competitive inhibition of Hpa1, and that, rather, Hpa2 has pleiotropic positive effects for the vasculature by regulating HS-dependent processes on the EC surface. In addition to the modulatory effects on growth-factor signaling addressed in this study, Hpa2 has the potential to regulate other HS-dependent processes including coagulatory and inflammatory cascades. Our work here establishes a solid foundation for future studies aimed at investigating that potential.

### Summary

In conclusion, this research establishes, for the first time, the role of endogenous Hpa2 as a systemic circulating molecule that maintains endothelial integrity and homeostasis. We show that Hpa2 derives primarily from endocrine organs and strongly binds HS, thereby controlling HS-dependent processes on the EC surface, including those mediated by Hpa1, VEGFA_165_, and FGF2. Our findings underscore the importance of the eGCX and its regulation for EC physiology and extend the spectrum of eGCX-targeting therapies for vascular disease. Hpa2 may serve as a pioneer for a novel class of protein- and peptide-based eGCX-stabilizing drugs.

## Acknowledgements

We thank Dr. Hans Bakker (Department of Biochemistry Hannover Medical School) for access and instructions to the chromatography devices. Lynne Staggs, Pat Schroder, Petra Wübbolt-Lehmann, and Birgit Habermeier for excellent technical support. The Hannover Medical School Research Core Unit for Laser Microscopy for continuous support. We would like to acknowledge the support from Chris Smith in the MDI Biological Laboratory (MDIBL) Sequencing Facility, Frederic Bonnet in the MDIBL Light Microscopy Facility and the whole team from the Animal Resources Core (Cores and Facilities supported by the Maine INBRE NIH P20GM103423 and the MDIBL COBRE NIH P20GM104318). Research reported in this publication was supported by an Institutional Development Award (IDeA) from the National Institute of General Medical Sciences of the National Institutes of Health under grant numbers P20GM103423 and P20GM104318. We are grateful to Stephen Sampson for critical editing the manuscript. Olivia Lentchner for designing the graphical model.

## Abbreviations

EC: Endothelial cell
ECM: Extracellular matrix
eGCX: Endothelial glycocalyx
ERK: Extracellular-signal-regulated kinase
FGF: Fibroblast growth factor
FGFR: FGF receptor
GAG: Glycosaminoglycan
GFB: Glomerular filtration barrier
hHpa2: Human heparanase 2
HMEC-1: Human microvascular endothelial cell 1
Hpa1: Heparanase
Hpa2: Heparanase 2
hpf: Hours post fertilization
HS: Heparan sulfate
ISV: Intersegmental vessel
LOF: Loss-of-function
MAPK: Mitogen-activated protein kinase
MEF: Maximum eye fluorescence
MO: Morpholino
mTOR: Mammalian target of rapamycin
PI3K: Phosphatidylinositol 3-kinases,
RTK: Receptor tyrosine kinase
VEGFA165: Vascular endothelial growth factor A165
VEGFR2: VEGF receptor 2

## References

1. Ricard N, Bailly S, Guignabert C, Simons M. The quiescent endothelium: signalling pathways regulating organ-specific endothelial normalcy. Nature Reviews Cardiology. 2021;18:565–580. 10.1038/s41569-021-00517-4

2. Reitsma S, Slaaf DW, Vink H, Van Zandvoort MAMJ, Oude Egbrink MGA. The endothelial glycocalyx: Composition, functions, and visualization. Pflugers Archive. 2007;454:345–359. doi: 10.1007/s00424-007-0212-8.

3. Weinbaum S, Cancel LM, Fu BM, Tarbell JM. The Glycocalyx and Its Role in Vascular Physiology and Vascular Related Diseases. Cardiovascular Engineering and Technology. 2021; 12:37–71. 10.1007/s13239-020-00485-9.

5. Florian JA, Kosky JR, Ainslie K, Pang Z, Dull RO, Tarbell JM. Heparan sulfate proteoglycan is a mechanosensor on endothelial cells. Circulation Research. 2003;93. 10.1161/01.RES.0000101744.47866.D5

4. Butler MJ, Down CJ, Foster RR, Satchell SC. The Pathological Relevance of Increased Endothelial Glycocalyx Permeability. American Journal of Pathology. 2020;190:742– 751. doi: 10.1016/j.ajpath.2019.11.015

6. Li W, Johnson DJD, Esmon CT, Huntington JA. Structure of the antithrombin-thrombin-heparin ternary complex reveals the antithrombotic mechanism of heparin. Nature Structural and Molecular Biology. 2004;11:857–862. 10.1038/nsmb811.

7. Mulivor AW, Lipowsky HH. Role of glycocalyx in leukocyte-endothelial cell adhesion. American Journal of Physiology. 2002; 283:1282–1291. 10.1152/ajpheart.00117.2002

8. Becker BF, Chappell D, Bruegger D, Annecke T, Jacob M. Therapeutic strategies targeting the endothelial glycocalyx: Acute deficits, but great potential. Cardiovascular Research. 2010;87:300–310. 10.1093/cvr/cvq137

9. Esko JD, Lindahl U. Molecular diversity of heparan sulfate. Journal of Clinical Investigation. 2001;108:169–173. doi: 10.1172/JCI13530.

10. Xu D, Esko JD. Demystifying heparan sulfate-protein interactions. Annual Review of Biochemistry. 2014;83:129–157. doi:10.1146/annurev-biochem-060713-035314

11. Schmidt EP, Yang Y, Janssen WJ, Gandjeva A, Perez MJ, Barthel L, Zemans RL, Bowman JC, Koyanagi DE, Yunt ZX et al. The pulmonary endothelial glycocalyx regulates neutrophil adhesion and lung injury during experimental sepsis. Nature Medicine. 2012;18:1217–1223. doi:10.1038/nm.2843

12. Chappell D, Jacob M, Hofmann-Kiefer K, Rehm M, Welsch U, Conzen P, Becker BF. Antithrombin reduces shedding of the endothelial glycocalyx following ischaemia/reperfusion. Cardiovascular Research. 2009;83:388–396. doi:10.1093/cvr/cvp097.

13. Nieuwdorp M, Van Haeften TW, Gouverneur MCLG, Mooij HL, Van Lieshout MHP, Levi M, Meijers JCM, Holleman F, Hoekstra JBL, Vink H et al. Loss of Endothelial Glycocalyx During Acute Hyperglycemia Coincides With Endothelial Dysfunction and Coagulation Activation In Vivo. Diabetes. 2006, 480–486. 10.2337/diabetes.55.02.06.db05-1103.

14. Banerjee S, Mwangi JG, Stanley TK, Mitra R, Ebong EE. Regeneration and Assessment of the Endothelial Glycocalyx to Address Cardiovascular Disease. Industrial & Engineering Chemical Research. 2021;60:17328–17347. 10.1021/acs.iecr.1c03074.

15. Vlodavsky I, Friedmann Y, Elkin M, Aingorn H, Atzmon R, Ishai-Michaeli R, Bitan M, Pappo O, Peretz T, Michael I et al.. Mammalian heparanase: Gene cloning, expression and function in tumor progression and metastasis. Nature Medicine 1999;5:793–802. 10.1038/10518.

16. Gil N, Goldberg R, Neuman T, Garsen M, Zcharia E, Rubinstein AM, Van Kuppevelt T, Meirovitz A, Pisano C, Li JP, Van Der Vlag J et al.. Heparanase is essential for the development of diabetic nephropathy in mice. Diabetes. 2012;61:208–216. 10.2337/db11-1024.

17. Nguyen TK, Paone S, Baxter AA, Mayfosh AJ, Phan TK, Chan E, Peter K, Poon IKH, Thomas SR, Hulett MD. Heparanase promotes the onset and progression of atherosclerosis in apolipoprotein E gene knockout mice. Atherosclerosis. 2024;392. 10.1016/j.atherosclerosis.2024.117519.

18. McKenzie E, Tyson K, Stamps A, Smith P, Turner P, Barry R, Hircock M, Patel S, Barry E, Stubberfield C et al.. Cloning and expression profiling of Hpa2, a novel mammalian heparanase family member. Biochemical and Biophysical Research Communication. 2000;276:1170–1177. 10.1006/bbrc.2000.3586

19. Levy-Adam F, Feld S, Cohen-Kaplan V, Shteingauz A, Gross M, Arvatz G, Naroditsky I, Ilan N, Doweck I, Vlodavsky I. Heparanase 2 interacts with heparan sulfate with high affinity and inhibits heparanase activity. Journal of Biological Chemistry. 2010;285:28010–28019. 10.1074/jbc.M110.116384.

20. Gross-Cohen M, Feld S, Arvatz G, Ilan N, Vlodavsky I. Elucidating the Consequences of Heparan Sulfate Binding by Heparanase 2. Frontiers in Oncology. 2021;10. doi:10.3389/fonc.2020.627463.

21. Liu J, Knani I, Gross-Cohen M, Hu J, Wang S, Tang L, Ilan N, Yang S, Vlodavsky I. Role of heparanase 2 (Hpa2) in gastric cancer. Neoplasia. 2021;23:966–978. 10.1016/j.neo.2021.07.010.

22. Kiyan Y, Tkachuk S, Kurselis K, Shushakova N, Stahl K, Dawodu D, Kiyan R, Chichkov B, Haller H. Heparanase-2 protects from LPS-mediated endothelial injury by inhibiting TLR4 signalling. Scientific Reports. 2019;9. doi:10.1038/s41598-019-50068-5.

23. Buijsers B, Garsen M, de Graaf M, Bakker-van Bebber M, Guo C, Li X, van der Vlag J. Heparanase-2 protein and peptides have a protective effect on experimental glomerulonephritis and diabetic nephropathy. Frontiers in Pharmacology. 2023;14. 10.3389/fphar.2023.1098184.

24. Kimmel CB, Ballard WW, Kimmel SR, Ullmann B, Schilling TF. Stages of embryonic development of the zebrafish. Developmental Dynamics. 1995;203:253–310. 10.1002/aja.1002030302.

25. Stuart HM, Roberts NA, Hilton EN, McKenzie EA, Daly SB, Hadfield KD, Rahal JS, Gardiner NJ, Tanley SW, Lewis MA, et al.. Urinary tract effects of HPSE2 mutations. Journal of the American Society of Nephrology. 2015;26:797–804. 10.1681/ASN.2013090961.

26. Guo C, Kaneko S, Sun Y, Huang Y, Vlodavsky I, Li X, Li ZR, Li X. A mouse model of urofacial syndrome with dysfunctional urination. Human Molecular Genetics. 2014;24:1991–1999. 10.1093/hmg/ddu613

27. Stahl K, Gronski PA, Kiyan Y, Seelinger B, Bertram A, Pape T, Welte T, Hoeper MM, Haller H, David S. Injury to the endothelial glycocalyx in critically Ill patients with COVID-19. American Journal of Respiratory and Critical Care Medicine. 2020;208:1178–1181. 10.1164/rccm.202007-2676LE

28. Stahl K, Hillebrand UC, Kiyan Y, Seeliger B, Schmidt JJ, Schenk H, Pape T, Schmidt BMW, Welte T, Hoeper MM et al.. Effects of therapeutic plasma exchange on the endothelial glycocalyx in septic shock. Intensive Care Medicine Experimental. 2021;9. 10.1186/s40635-021-00417-4.

29. Roberts NA, Woolf AS, Stuart HM, Thuret R, McKenzie EA, Newman WG, Hilton EN. Heparanase 2, Mutated in urofacial syndrome, Mediates peripheral neural development in Xenopus. Human Molecular Genetics. 2014;23:4302–4314. 10.1093/hmg/ddu147.

30. Xie J, Farage E, Sugimoto M, Anand-Apte B. A novel transgenic zebrafish model for blood-brain and blood-retinal barrier development. BMC Developmental Biology. 2010. 10.1186/1471-213X-10-76.

31. Singh A, Satchell SC, Neal CR, McKenzie EA, Tooke JE, Mathieson PW. Glomerular endothelial glycocalyx constitutes a barrier to protein permeability. Journal of the American Society of Nephrology. 2007;18:2885–2893. 10.1681/ASN.2007010119.

32. Mottarella SE, Beglov D, Beglova N, Nugent MA, Kozakov D, Vajda S. Docking server for the identification of heparin binding sites on proteins. J Chem Inf Model. 2014; 54:2068–2078. 10.1021/ci500115j.

33. Corti F, Wang Y, Rhodes JM, Atri D, Archer-Hartmann S, Zhang J, Zhuang ZW, Chen D, Wang T, Wang Z, Azadi P, Simons M. N-terminal syndecan-2 domain selectively enhances 6-O heparan sulfate chains sulfation and promotes VEGFA165-dependent neovascularization. Nature Communication. 2019;10. 10.1038/s41467-019-09605-z.

34. Yang X, Liaw L, Prudovsky I, Brooks PC, Vary C, Oxburgh L, Friesel R. Fibroblast Growth Factor Signaling in the Vasculature. Current Atherosclerosis Reports. 2015;17. 10.1007/s11883-015-0509-6.

35. Simons M, Gordon E, Claesson-Welsh L. Mechanisms and regulation of endothelial VEGF receptor signalling. Nature Reviews Molecular Cell Biology. 2016; 17:611–625. 10.1038/nrm.2016.87

36. Schlessinger J, Plotnikov AN, Ibrahimi OA, Eliseenkova A V, Yeh BK, Yayon A, Linhardt RJ, Mohammadi M. Crystal Structure of a Ternary FGF-FGFR-Heparin Complex Reveals a Dual Role for Heparin in FGFR Binding and Dimerization. Molecular Cell; 2000: 1–743–750. 10.1016/S1097-2765(00)00073-3.

37. Ferrara N, Davis-Smyth T. The Biology of Vascular Endothelial Growth Factor. Endocrine Reviews; 1997. 10.1210/edrv.18.1.0287

38. Vlodavsky I, Hilwi M, Kayal Y, Soboh S, Ilan N. Impact of heparanase-2 (Hpa2) on cancer and inflammation: Advances and paradigms. FASEB Journal. 2024;38. 10.1096/fj.202400286R.

39. Knani I, Singh P, Gross-Cohen M, Aviram S, Ilan N, Sanderson RD, Aronheim A, Vlodavsky I. Induction of heparanase 2 (Hpa2) expression by stress is mediated by ATF3. Matrix Biology. 2022;105:17–30. 10.1016/j.matbio.2021.11.001.

40. Hu S, Zhao X, Li R, Hu C, Wu H, Li J, Zhang Y, Xu Y. Activating transcription factor 3, glucolipid metabolism, and metabolic diseases. Journal of Molecular Cell Biology. 2022;14. 10.1093/jmcb/mjac067.

41. Pang J, Zhang S, Yang P, Hawkins-Lee B, Zhong J, Zhang Y, Ochoa B, Agundez JAG, Voelckel MA, Gu W, et al.. Loss-of-Function Mutations in HPSE2 Cause the Autosomal Recessive Urofacial Syndrome. The American Journal of Human Genetics. 2010;86:957–962. 10.1016/j.ajhg.2010.04.016

42. Daly SB, Urquhart JE, Hilton E, McKenzie EA, Kammerer RA, Lewis M, Kerr B, Stuart H, Donnai D, Long DA et al.. Mutations in HPSE2 Cause Urofacial Syndrome. The American Journal of Human Genetics. 2010;86:963–969. 10.1016/j.ajhg.2010.05.006.

43. Manak I, Gurney AM, McCloskey KD, Woolf AS, Roberts NA. Dysfunctional bladder neurophysiology in urofacial syndrome Hpse2 mutant mice. Neurourology Urodynamics. 2020;39:1930–1938. 10.1002/nau.24450.

44. Kayal Y, Barash U, Naroditsky I, Ilan N, Vlodavsky I. Heparanase 2 (Hpa2)- a new player essential for pancreatic acinar cell differentiation. Cell Death Disease. 2023;14. 10.1038/s41419-023-05990-y.

45. Lee S, Chen TT, Barber CL, Jordan MC, Murdock J, Desai S, Ferrara N, Nagy A, Roos KP, Iruela-Arispe ML. Autocrine VEGF Signaling Is Required for Vascular Homeostasis. Cell. 2007;130:691–703. 10.1016/j.cell.2007.06.054.

46. Miquerol L, Langile L, Nagy A. Embryonic development is disrupted by modest increases in vascular endothelial growth factor gene expression. Development. 2000;127:3941–3946. 10.1242/dev.127.18.3941.

47. Ruhrberg C, Gerhardt H, Golding M, Watson R, Ioannidou S, Fujisawa H, Betsholtz C, Shima DT. Spatially restricted patterning cues provided by heparin-binding VEGF-A control blood vessel branching morphogenesis. Genes Dev. 2002 Oct 15;16(20):2684–98. 10.1101/gad.242002.

48. Jakobsson L, Kreuger J, Holmborn K, Lundin L, Eriksson I, Kjellén L, Claesson-Welsh L. Heparan Sulfate in trans Potentiates VEGFR-Mediated Angiogenesis. Developmental Cell. 2006;10:625–634. 10.1016/j.devcel.2006.03.009.

49. Préau L, Lischke A, Merkel M, Oegel N, Weissenbruch M, Michael A, Park H, Gradl D, Kupatt C, le Noble F. Parenchymal cues define Vegfa-driven venous angiogenesis by activating a sprouting competent venous endothelial subtype. Nature Communications. 2024;15. 10.1038/s41467-024-47434-x.

50. Senger DR, Galli SJ, Dvorak AM, Perruzi C, Harvey S V., Dvorak HF. Tumor Cells Secrete a Vascular Permeability Factor That Promotes Accumulation Of Ascites Fluid. Science. 1983;219:983–985. 10.1126/science.6823562.

51. Wild R, Klems A, Takamiya M, Hayashi Y, Strähle U, Ando K, Mochizuki N, Van Impel A, Schulte-Merker S, Krueger J, Preau L, Le Noble F. Neuronal sFlt1 and Vegfaa determine venous sprouting and spinal cord vascularization. Nature Communications. 2017;8. 10.1038/ncomms13991.

52. Eremina V, Sood M, Haigh J, Nagy A, Lajoie G, Ferrara N, Gerber HP, Kikkawa Y, Miner JH, Quaggin SE. Glomerular-specific alterations of VEGF-A expression lead to distinct congenital and acquired renal diseases. J Clin Invest. 2003; 111(5):707–16. doi: 10.1172/JCI17423.

53. Murakami M, Nguyen LT, Zhang ZW, Moodie KL, Carmeliet P, Stan R V., Simons M. The FGF system has a key role in regulating vascular integrity. Journal of Clinical Investigation. 2008;118:3355–3366. 10.1172/JCI35298.

54. Elkin M, Ilan N, Ishai-Michaeli R, Friedmann Y, Papo O, Pecker I, Vlodavsky I. Heparanase as mediator of angiogenesis: mode of action. FASEB Journal. 2001;15:1661–1663. 10.1096/fj.00-0895fje

55. Ricard N, Scott RP, Booth CJ, Velazquez H, Cilfone NA, Baylon JL, Gulcher JR, Quaggin SE, Chittenden TW, Simons M. Endothelial ERK1/2 signaling maintains integrity of the quiescent endothelium. Journal of Experimental Medicine. 2019;216:1874–1890. 10.1084/jem.20182151.

56. Boels MGS, Avramut MC, Koudijs A, Dane MJC, Lee DH, Van Der Vlag J, Koster AJ, Van Zonneveld AJ, Van Faassen E, Gröne HJ. et al.. Atrasentan reduces albuminuria by restoring the glomerular endothelial glycocalyx barrier in diabetic nephropathy. Diabetes. 2016; 65:2429–2439. 10.2337/db15-1413.

57. Crompton M, Ferguson JK, Ramnath RD, Onions KL, Ogier AS, Gamez M, Down CJ, Skinner L, Wong KH, Dixon LK et al.. Mineralocorticoid receptor antagonism in diabetes reduces albuminuria by preserving the glomerular endothelial glycocalyx. JCI Insight. 2023;8. 10.1172/jci.insight.154164.

58. Gamez M, Elhegni HE, Fawaz S, Ho KH, Campbell NW, Copland DA, Onions KL, Butler MJ, Wasson EJ, Crompton M et al.. Heparanase inhibition as a systemic approach to protect the endothelial glycocalyx and prevent microvascular complications in diabetes. Cardiovascular Diabetology. 2024;23. 10.1186/s12933-024-02133-1.

59. Song JW, Zullo JA, Liveris D, Dragovich M, Zhang XF, Goligorsky MS. Therapeutic restoration of endothelial glycocalyx in sepsis. Journal of Pharmacology and Experimental Therapeutics. 2017;361:115–121. 10.1124/jpet.116.239509.

60. De Boer C, Armstrong Z, Lit VAJ, Barash U, Ruijgrok G, Boyango I, Weitzenberg MM, Schröder SP, Sarris AJC, Meeuwenoord NJ, Bule P, Kayal Y, Ilan N, Cod JDC, Vlodavsky I, Overkleeft HS, Davies GJ, Wu L. Mechanism-based heparanase inhibitors reduce cancer metastasis in vivo. PNAS. 2022;119. 10.1073/pnas.

61. Heyman B, Yang Y. Mechanisms of heparanase inhibitors in cancer therapy. Experimental Hematology. 2016;44:1002–1012. 10.1016/j.exphem.2016.08.006.

62. Ferrara, N., Adamis, A. Ten years of anti-vascular endothelial growth factor therapy. Nat Rev Drug Discov 15, 385–403 (2016). 10.1038/nrd.2015.17.

63. Lawson ND, Weinstein BM. In vivo imaging of embryonic vascular development using transgenic zebrafish. Developmental Biology. 2002;248:307–318. 10.1006/dbio.2002.0711.

64. Wang Y, Kaiser MS, Larson JD, Nasevicius A, Clark KJ, Wadman SA, Roberg-Perez SE, Ekker SC, Hackett PB, McGrail M, Essner JJ. Moesin1 and Ve-cadherin are required in endothelial cells during in vivo tubulogenesis. Development. 2010;137:3119– 3128. 10.1242/dev.048785.

65. Westerfield, M., 1993. The Zebrafish Book: a Guide for the Laboratory Use of Zebrafish (Brachydanio Rerio).

66. Thisse B, Thisse C. In situ hybridization on whole-mount zebrafish embryos and young larvae. Methods in Molecular Biology. 2014;1211:53–67.

67. McDowell KP, Berthiaume AA, Tieu T, Hartmann DA, Shih AY. VasoMetrics: Unbiased spatiotemporal analysis of microvascular diameter in multi-photon imaging applications. Quantitative Imaging in Medicine and Surgery. 2021;11:969–982. 10.21037/qims-20-920.

68. Jörns A, Wedekind D, Jähne J, Lenzen S. Pancreas pathology of latent autoimmune diabetes in adults (LADA) in patients and in a LADA rat model compared with type 1 diabetes. Diabetes. 2020;69:624–633. 10.2337/db19-0865.

69. Pende M, Vadiwala K, Schmidbaur H, Stockinger AW, Murawala P, Saghafi S, S Dekens MP, Becker K, Revilla-i-Domingo R, Papadopoulos SC. A versatile depigmentation, clearing, and labeling method for exploring nervous system diversity. Science Advances. 2020;6. 10.1126/sciadv.aba0365.

70. Schenk H, Müller-Deile J, Schroder P, Bolaños-Palmieri P, Beverly-Staggs L, White R, Bräsen JH, Haller H, Schiffer M. Characterizing renal involvement in Hermansky-Pudlak Syndrome in a zebrafish model. Scientific Reports. 2019;9. 10.1038/s41598-019-54058-5.

71. Wiznerowicz M, Trono D. Conditional Suppression of Cellular Genes: Lentivirus Vector-Mediated Drug-Inducible RNA Interference. Journal of Virology. 2003;77:8957–8961. 10.1128/JVI.77.16.8957-8961.2003

